# Quantum biological insights into CRISPR-Cas9 sgRNA efficiency from explainable-AI driven feature engineering

**DOI:** 10.1101/2022.06.03.494572

**Authors:** Jaclyn M. Noshay, Tyler Walker, Jonathon Romero, Erica Prates, Carrie Eckert, Stephan Irle, David Kainer, Daniel A. Jacobson

**Author notes:** This manuscript has been authored by UT-Battelle, LLC under Contract No. DE-AC05-00OR22725 with the U.S. Department of Energy. The United States Government retains and the publisher, by accepting the article for publication, acknowledges that the United States Government retains a non-exclusive, paid-up, irrevocable, world-wide license to publish or reproduce the published form of this manuscript, or allow others to do so, for United States Government purposes. The Department of Energy will provide public access to these results of federally sponsored research in accordance with the DOE Public Access Plan (http://energy.gov/downloads/doe-public-access-plan).

## Abstract

CRISPR-Cas9 tools have transformed genetic manipulation capabilities in the laboratory. Empirical rules-of-thumb have been established for only a narrow range of model organisms, and mechanistic underpinnings for sgRNA efficiency remain poorly understood. This work establishes a novel feature set and new public resource, produced with quantum chemical tensors, for interpreting and predicting sgRNA efficiency. Feature engineering for sgRNA efficiency is performed using an explainable-artificial intelligence model; iterative Random Forest (iRF). By encoding quantitative attributes of position-specific sequences for *E. coli* sgRNAs, we identify important traits for sgRNA design in bacterial species. Additionally, we show that expanding positional encoding to quantum descriptors of base-pair, dimer, trimer and tetramer sequences captures intricate interactions in local and neighboring nucleotides of the target DNA. These features highlight variation in CRISPR-Cas9 sgRNA dynamics between *E. coli* and *H. sapien* genomes. These novel encodings of sgRNAs greatly enhance our understanding of the elaborate quantum biological processes involved in CRISPR-Cas9 machinery.

---

CRISPR-Cas9 is revolutionizing genome-editing through the use of a single guide RNA (sgRNA) to direct precise cleavage at endogenous locations in the genome ^1,2^. The first step to engineer or modify a specific region using CRISPR-Cas9 is to computationally predict cutting efficiencies of potential sgRNAs. The CRISPR-Cas9 system depends on this designed sgRNA to target the protein complex to a region flanked by a 3’NGG protospacer adjacent motif (PAM). The CRISPR-Cas9 system is only successful if both specificity and efficiency occur at the target loci 3. To inform sgRNA sequence choices, genomic feature analyses have associated sgRNA attributes with cutting efficiency for CRISPR-Cas9 systems ^4–8^.

Predicting sgRNA efficiency requires careful consideration of relationships among the sgRNA sequence, genomic features of the target region, and activity within the CRISPR-Cas9 system. Some of these relationships have been studied extensively 9. Among them, nucleotide composition of the target sequence is the most thoroughly studied contributor to sgRNA efficiency ^3,10–12^. Specific nucleotide patterns have been associated with sgRNA efficiency; including the presence of guanine and absence of thymine near the PAM sequence, preference for cytosine near the cut site, and overall GC content ^3,11,13^. The seed region – defined as the five to ten bases of the target sequence nearest the PAM – is of central consideration for these patterns in sgRNA sequence composition ^10,12,14^.

While nucleotide sequence patterns are observed across species, their influence on integration and cleavage with CRISPR-Cas9 may vary ^1,15–18^. The added complexities of chromatin structure have started to be considered, enhancing understanding of CRISPR-Cas9 dynamics. For example, human models were expanded with information about the insertion point within the gene sequence ^19^ and secondary structure of the target sequence ^20,21^. Target regions with low nucleosome occupancy and high chromatin accessibility have also been investigated ^22–26^. These structural nuances underscore even greater variation in the CRISPR-Cas9 system across organisms.

DNA is less protected in prokaryotic cells than in eukaryotic cells because of a simpler chromatin structure; and target regions are often more accessible 3. In contrast, mammalian cells have highly-active non-homologous end-joining systems, which induce repair mechanisms for the DNA double strand break during CRISPR-Cas9 integration. In prokaryotes, sgRNA activity is correlated with cellular survival because double stranded breaks are lethal to the cell ^27^. These pronounced differences in structure and function illuminate, in part, why models trained for mammalian species have failed to provide sgRNAs that consistently integrate with the target sequence in other kingdoms. This insufficiency spurred development of organism-tailored models, including those for plants 1, yeast ^18^ and bacteria ^28^. Expanding the breadth and chemical specificity of model feature sets provides useful avenues for extending state-of-the-art sgRNA efficiency prediction to other organisms and non-model species. To achieve this next level of model prediction power, quantum chemical properties warrant consideration.

Bridging chemistry and physics, quantum chemical properties capture the ways in which electron density impact the reactivities and energetics of molecules. Some properties, such as the HOMO-LUMO gap (highest occupied molecular orbital-lowest unoccupied molecular orbital energy gap), describe how electron density is distributed among atoms. Meanwhile, other properties, like hydrogen-bonding energy or π-stacking energy, describe how a system’s total energy changes as molecules interact. Such properties depend on how the molecular electron densities shift in response to one another. Incorporating quantum chemical detail when characterizing or predicting biological processes has been transformative for biology; providing new frameworks for viewing processes, identifying novel features, and enhancing mechanistic understandings ^29,30^. This work spotlights quantum properties including HOMO-LUMO gaps, hydrogen bonding, and stacking interactions to investigate the complex molecular interactions of the DNA double helix and the DNA CRISPR-Cas9 sgRNA hybrid.

Machine learning models excel at identifying patterns in data to inform outcomes; but the power behind these algorithms is bottlenecked by the depth and breadth of the training data. Current methods of feature evaluation for CRISPR-Cas9 efficiency are trained on experimental sgRNA cutting efficiency data from a narrow range of eukaryotic species, including human, mouse, and zebrafish 4. While these models’ species-by-species rules for sgRNA prediction are informative, their insights are rarely generalizable. Therefore, to develop advanced predictive models, the training data must be sufficiently detailed to capture the complexities of genomic structure and content that influence efficiencies of CRISPR-Cas9 integration and cleavage across systems.

Here we use machine learning approaches to unravel these species-dependent rules of sgRNA efficiency. Many current AI model generation approaches use techniques such as neural networks that can obscure associations behind a “black box” of decision schemes. We sought to understand feature contributions to cutting efficiency for *E.coli* through an explainable-artificial intelligence (XAI) approach. We used iterative Random Forest (iRF), an XAI method designed for model transparency and feature evaluation, to assess CRISPR-Cas9 efficiency and improve our understanding of the system’s underlying biological mechanisms. When trained on detailed feature sets, XAI models provide a shared basis for predicting sgRNA efficiency across organisms. This work extends sgRNA efficiency modeling to assess both *E. coli* and *H. sapien* datasets. Additionally, our model integrates a novel and interdisciplinary feature set that includes quantum chemical properties.

## Results

### Feature importance with iterative Random Forest

We assess iRF-based predictive models that leverage quantum descriptors for multiple degrees of base-pair polymerization. This data captures interactions within the sgRNA nucleotide sequence, along with properties of the individual bases. This approach combines the increased interpretability of XAI methods for feature interpretation with the novel incorporation of quantum chemical properties to further mechanistic understandings of CRISPR-Cas9. Models were generated for sgRNA efficiency in *E. coli* and *H. sapien*. Additionally, the variations in feature importance across kingdoms were assessed.

A publicly-available *E. coli* dataset was used to generate a detailed feature matrix to determine highly-influential properties for predicting CRISPR-Cas9 cutting efficiencies (Figure 1). In this model, the dependent variable (Y-vector) is the experimental cutting efficiency score for every sgRNA. We used quantum chemical features of nucleotides to capture the intricacies of the multi-step CRISPR-Cas9 mechanism. In addition, base-pair oligomers (kmers) up to tetramers were incorporated to enhance understanding of nucleotide position within the sgRNA structure through binary encoding (one-hot kmers) and quantum chemical properties (quantum kmers). Features previously determined to influence CRISPR-Cas9 efficiency in mammalian species were also encoded; including GC content, melting temperature, and one-hot encoding of single and paired nucleotides of the target sequence. In addition, we included quantitative measures such as the distance between the target sequence and the nearest PAM (NGG sequence) and location of the target sequence relative to the nearest gene. To sharpen insight into the particular contributions of each feature, predictive capabilities were assessed by Pearson correlations and accuracy (R^2^) metrics (Supplemental Results). The iRF model using the complete feature matrix results in a predictive accuracy of 0.51, which is comparable with the most predictive models currently available. Quantum kmers and one-hot kmers contribute the largest feature set contributions, pinpointing isolated features important for sgRNA engineering.

**Figure 1:**
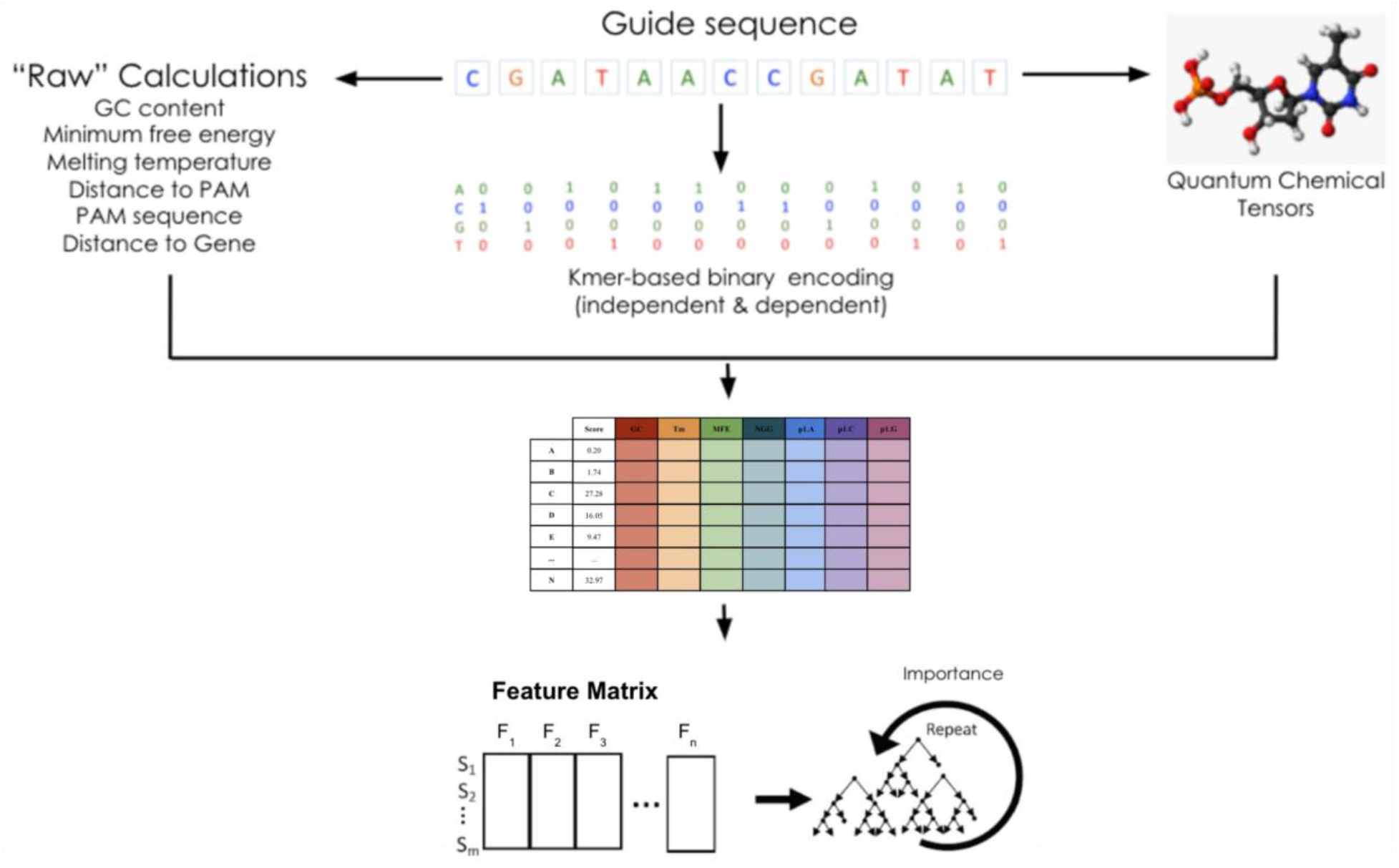
Explainable-AI method for analysis of feature importance on prediction of sgRNA efficiency. Features are formatted to generate a wide matrix with rows representing each sgRNA, the corresponding experimental cutting efficiency, and columns for all respective numeric feature values. This information is fed into an iterative Random Forest (iRF) analysis.

### Feature engineering highlights the role of quantum mechanics

A feature engineering approach enhances the understanding of factors influencing sgRNA efficiency by identifying the model’s most important variables. A total of 6,232 features capturing information in *E. coli* were used in an iRF model (the full *E. coli* feature set). This model was trained on 32,374 sgRNA and tested on 8,094 sgRNA sequences. This complete feature matrix cast as an iRF framework resulted in significant correlations between predicted and experimental sgRNA cutting efficiency values (Figure 2A). Furthermore, high prediction levels were found in the iRF model using only the quantum chemical properties feature set; and accuracy increased incrementally as additional features and kmers were incorporated (Figure 2A; Figure S1; Supplemental results). Below, we focus on a subset of features that contributed large effects to predicting sgRNA efficiency (Figure 2B-C).

**Figure 2:**
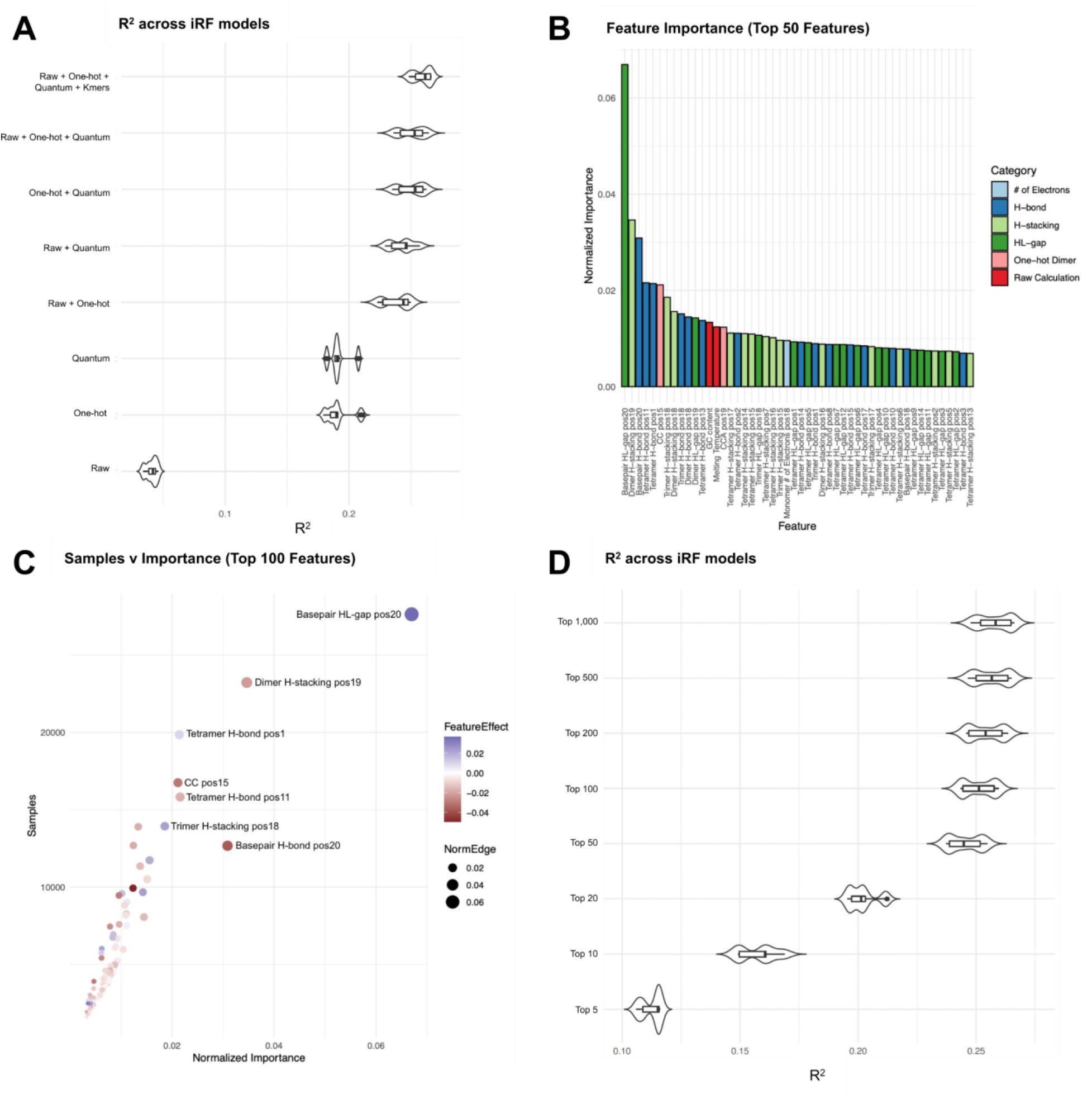
Identifying model variation based on feature input and assessing feature importance in E.coli. A) Violin plot of R squared values based on iRF model generation with isolated feature input (feature categories described in Table SX). B) The top 50 features from the full feature matrix iRF model ranked by normalized feature importance score and colored by feature category. C) Dot plot of features from full feature matrix iRF model showing the number of samples (sgRNAs) that were influenced by that feature (y-axis) versus the normalized importance of the feature (x-axis). Color is determined by the feature effect score (negative = red, positive = blue) and dot size is determined by the normalized importance score. D) Violin plot of R squared values for the top 5, 10, 20, 50, 100, 200, 500, and 1000 features based on full feature iRF model output showing the plateau of information gained from features beyond a certain level of importance.

Based on the feature importance values produced by iRF, top features emphasized positionally-encoded kmers of quantum chemical properties and one-hot encoding of the target sequence (Figure 2B). Top features were localized to positions 18 through 20 of the target sequence. This region is proximal to the sgRNA tailpin structure, the target DNA PAM sequence, and cut site for the Cas9 nuclease. The most important feature was the HOMO-LUMO energy gap for the base pair at position 20 of the target sequence (Figure 2B). This feature alone accounted for more than 6% of the variance in empirical sgRNA efficiency. The next most important feature was the base-pair dimer stacking energy at bases 19 and 20; accounting for ~3% of the variance. Hydrogen bond energy of the base pair at position 20 shows a similar contribution. Following these features in importance, we observe several position-dependent base pair dimer, trimer, and tetramer quantum chemical-encoded values. Each of these features accounted for 1-2% of the dataset variance. Several one-hot encoding sequences were also important features, including cytosine positions 15 and 16 (CC pos15) and a CCA beginning at position 19. Additionally, several features with high feature importance scores are consistent with trends in the literature, including GC content and melting temperature ^31^.

Each decision tree within an iRF model selects features based on their contribution to the predictive ability of the experimental dataset. Because of complex relationships between features and sgRNA, however, individual features may not be influential for every sgRNA in the model. Therefore, each feature’s average number of affected sgRNAs was also calculated for all decision trees within the iRF model. This average was compared with the total number of training data sgRNAs to determine the relative proportion of sgRNAs that each feature influenced (Figure 2C). The twenty highest-magnitude features affected between 20% and 85% of sgRNA samples. The three most-commonly influential features included the base pair 20 HOMO-LUMO gap energy, the base-pair 19-20 dimer hydrogen bond energy, and the base-pair 1-4 tetramer hydrogen bond energy (Figure 2C). These top features span both positive and negative associations with predicted cutting efficiency scores.

### Assessing feature association with sgRNA efficiency

Each feature was assigned a direction (positive or negative) and effect size, calculated with a random intersection tree (RIT)-based approach (Figure 3A; ^32^). These components describe a feature’s relationship with the cutting efficiency score, allowing for greater interpretation of that feature’s role in the CRISPR-Cas9 mechanism. For example, it has been shown that higher melting temperatures and greater GC content decrease guide efficiency ^31^. This anti-correlated relationship is demonstrated in our model by a negative feature effect value (Figure 3A/C). Important features exhibited both positive and negative correlations with the predicted cutting efficiency score.

**Figure 3:**
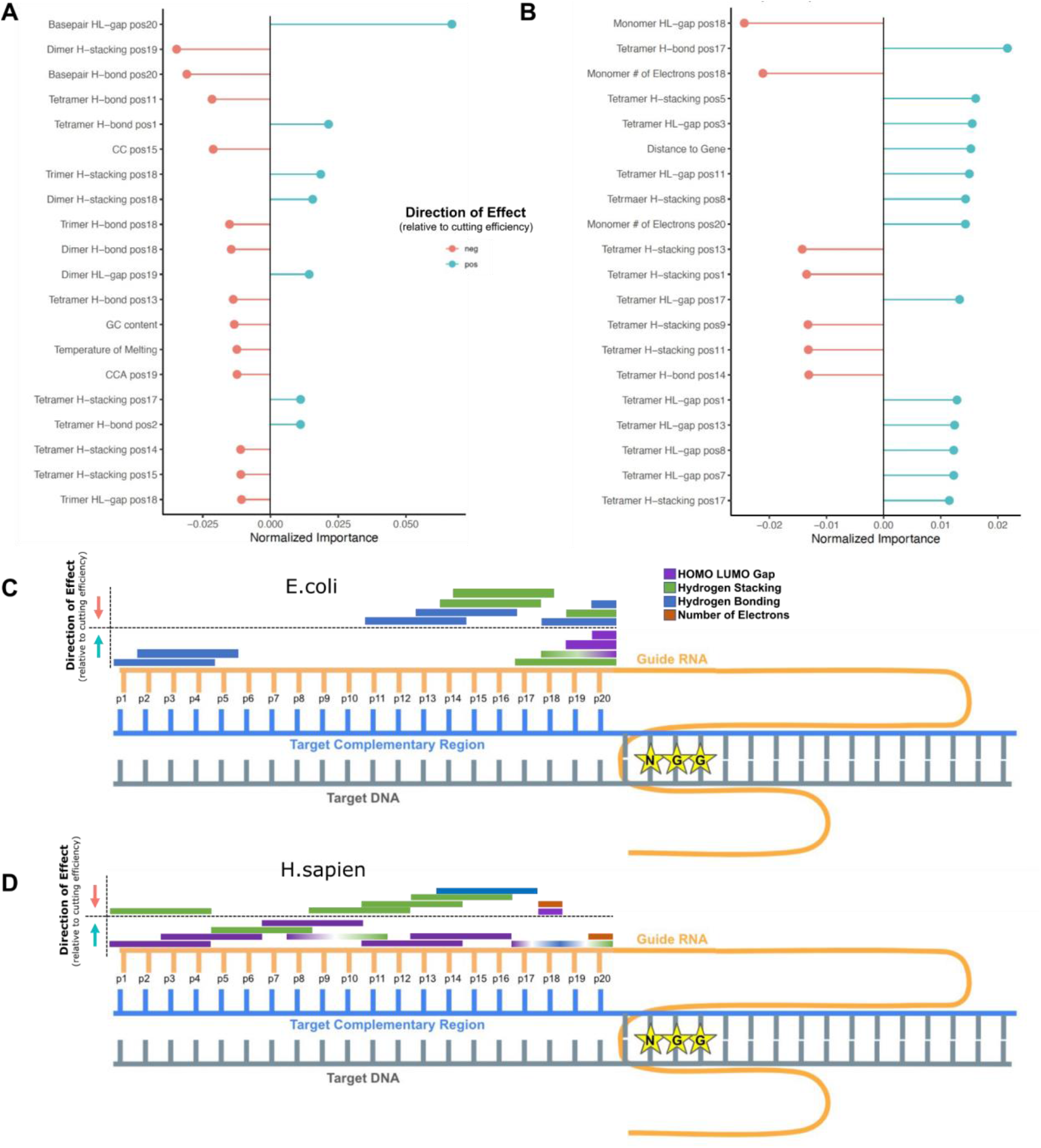
Explainable-AI interpretation through iRF output metrics and directional influence on cutting efficiency. A-B) The top 20 features from the full E.coli matrix ranked by normalized importance score and colored by the direction of the effect as positively correlated with the cutting efficiency score (blue) or anti-correlated with cutting efficiency score (pink) for *E.coli* (A) and *H.sapien* (B). C-D) sgRNA-DNA interaction highlighting quantum chemical property features of top importance and their localization for positively correlated association and anticorrelated associations with cutting efficiency scores in *E.coli* (C) and *H.sapien* (D). DNA strand represented in gray (target sequence) and blue (target complementary sequence), sgRNA shown in yellow, and PAM sequence displayed with NGG stars. Direction of feature effect is indicated with arrows, up (blue arrow) indicates a positively correlated relationship between the feature value and the cutting efficiency value. Feature bars indicate specific quantum properties (HL gap = purple, Stacking interactions = green, H-bonding = blue) and the length of the bar indicates the kmer size (multi-colored bars indicate the same k-mer at the same position has multiple features determined to be of high importance). The *E. coli* (C) model shows extensive localization with important features, primarily long kmers, focused in positions 11-20, with hydrogen bonding having outlier importance at position 1-5. Hydrogen bonding and stacking energy are observed to have both correlated and anti-correlated relationships with cutting efficiency (depending on their kmer and position) while HL-gap is consistent to a positive relationship closest to the PAM sequence. The *H. sapien* (D) model shows a greater continuum of location, many features stacked in positions 5-15) and feature-specific directional effect (hydrogen bonding, stacking energy, and HL-gap all found to have both positive and negative relationships with cutting efficiency dependent on the feature length and position) for features of high importance, but maintains that long kmers appear to be the most predictive. The number of electrons appears novel to *H. sapien* in comparison to *E. coli* when focused on top features of importance in the iRF model.

The top positional encoding features also showed varied directions of correlation with sgRNA cutting efficiency. Two essential features are the HOMO-LUMO gap and hydrogen bond strength at position 20 of the sgRNA and target sequence (Figure 3A/C, S2A-B). The HOMO-LUMO gap is positively correlated with sgRNA cutting efficiency, while the hydrogen bond strength at the same position is anti-correlated. Further, the directionality of the hydrogen bond strength effect varies by position and encoding length – whether base-pair monomers, dimers, trimers, or tetramers are considered. Hydrogen bond strength in positions 18-20 have negative effects, while hydrogen bond strength at position 1 has a positive effect. This contrast indicates varied preferences for hydrogen bonding energy across regions of the target sequence. One-hot encoding indicates position 15 CC as anti-correlated, while position 19 GC is positively correlated with sgRNA cutting efficiency. Additionally, our model indicates that increased distance to PAM is anti-correlated with sgRNA cutting efficiency.

### Quantum chemical insights into kingdom-specific dynamics

Current species-trained models in the literature are inadequate for prediction across organisms. To assess organism specificity of our iRF model, we tested the efficacy of the full *E. coli*-trained model on several publicly accessible *H. sapien* datasets ^10,33^. The resulting predictions were extremely poor, with a Pearson correlation of 0.016. This reinforces the idea that features determined by models trained on experimental data from a single species are not predictive across species, particularly where varied CRISPR-Cas9 interactions and complex DNA structures contribute.

*E. coli* and *H. sapien* represent different kingdoms; Eubacteria and Animalia. These classifications span single-celled to multi-celled organisms; varied organellar makeup and diversity in genomic and epigenomic structures and compositions. To compare the predictive capability of the newly integrated feature set across kingdoms, we generated a model trained on *H.sapien*-specific data ^10,33^. The full feature matrix was generated as described for the *E. coli* model, using the specified sgRNA sequences in the Doench et al. 2014 (1,278 sgRNAs) and Chuai et al. 2018 (16,749 sgRNAs) datasets. The iRF model was prepared with the same five-fold cross validation scheme as for the *E. coli* models. The resulting model had a Pearson correlation of 0.50 (Table 1). This is competitive with several of the top predictive cutting efficiency models currently available for human genome editing 4.

**Table 1:**
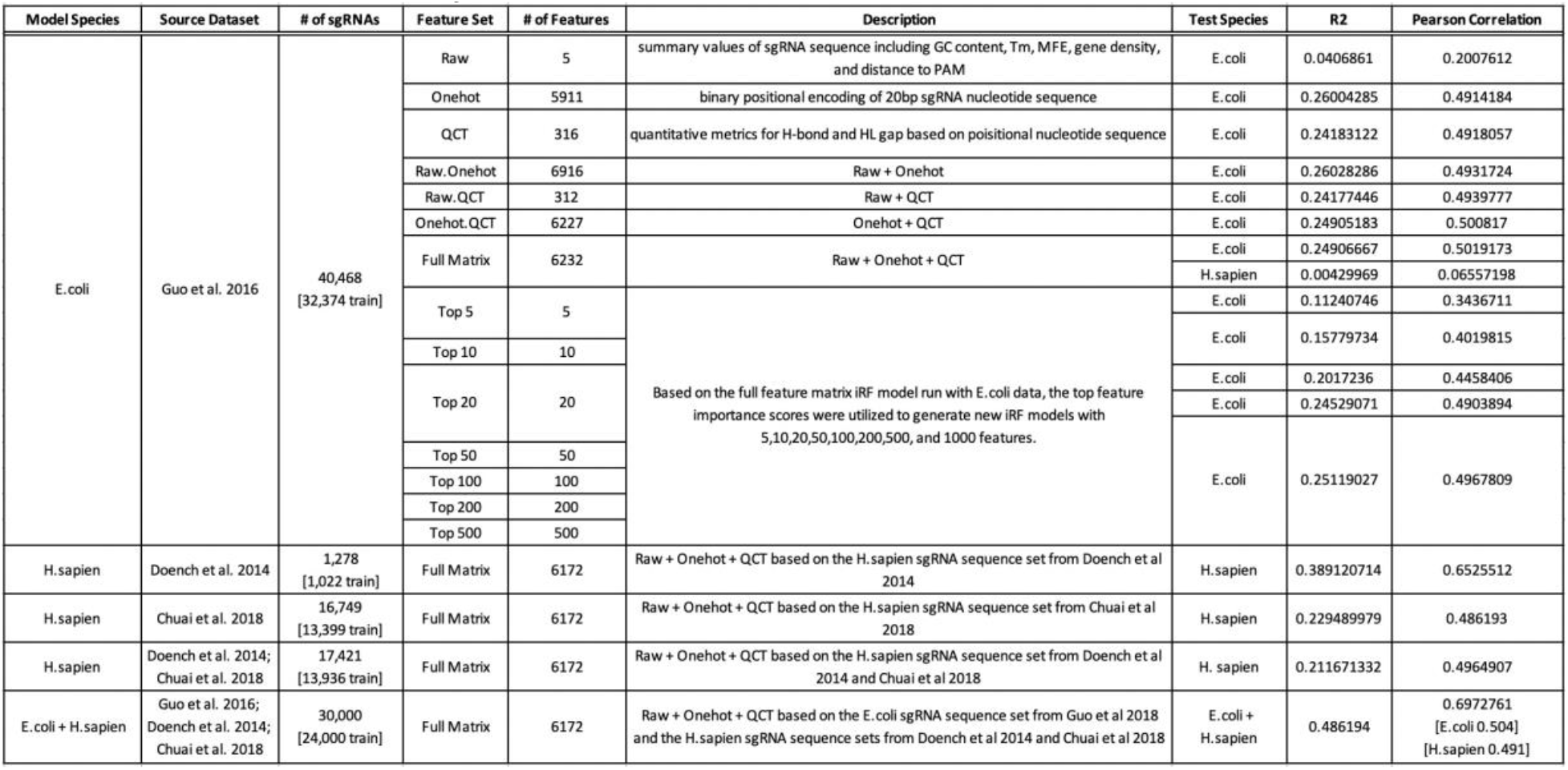
iRF model summary and metrics

To explore this result, we cross-examined the twenty highest-scoring features for each species-trained model (Figure 3). Similarly to those identified in the *E. coli* model, quantum chemical tensors in the target sequence’s seed region (sgRNA position 10-20, closest to the PAM sequence) appear to drive the *H. sapien* model prediction (Figure 3B). While quantum chemical tensors as a feature set are important for sgRNA efficiency in both *E. coli* and *H. sapien*, the particular features of importance vary considerably between models. In *E. coli*, the occupancy of frontier orbitals for PAM-adjacent nucleotides was a driving factor in CRISPR-Cas9 cutting (Figure 3C). In the *H. sapien* model, however, properties in central regions of the target sequence were highlighted, specifically positions 5-15 (Figure 3D). Key features for this model emphasize hydrogen bond energy and stacking interaction along with electron occupancy (Figure 3B/D). These features signpost novel mechanistic interpretations focused on central regions of the target sequence that can be explored in future biological studies.

## Discussion

Current sgRNA efficiency prediction models are limited by a narrow range of species data for training. To enhance the predictive accuracy, many models use deep learning techniques that can obscure interpretability of feature influence. This work sought to expand understanding of the factors influencing sgRNA efficiency through a bacterial dataset and an explainable-AI method. Furthermore, incorporation of quantum chemical properties provided novel insights into the interpretation of sgRNA efficiency dynamics and the CRISPR-Cas9 mechanism.

A panel of *E.coli* sgRNA sequences were encoded into a matrix incorporating PAM sequence distance, sgRNA melting temperature, GC content, one-hot binary encoding, and quantum chemical property encoding. This detailed feature set was used to train an XAI iRF model. The predictive capability of the machine learning model was enhanced by advanced kmer features (binary and quantum) and is competitive with currently-available models. The XAI methodology permitted investigation of the underlying features by quantifying feature importance scores.

Quantum chemical properties carried the highest importance for prediction of sgRNA efficiency. This feature set is novel in the domain of CRISPR-Cas9 models and enhances the model’s biological interpretability. Beyond position-specific sequence information, which is commonly encoded in a binary matrix, quantum chemical properties signpost the varied nucleotide interactions that mediate the CRISPR-Cas9 mechanism. The sgRNA seed region featured quantum properties with high predictive capacities. Descriptors of hydrogen bond energy, stacking interactions, and HOMO LUMO gaps enrich the interpretation of why this region plays a vital role in CRISPR-Cas9 efficiency. Particularly, we note indications of mechanistic competition for preferred structural features. We focus on three main themes: locality in the “seed region”, degree of base-pair polymerization, and mechanistic competition (Figure 3C).

The “seed” region, the five to ten base pairs on the target sequence’s 3’ end nearest to the PAM sequence and cleavage site, has been a focus of sgRNA construction across mammalian species. Several of the top *E. coli*-based features, specifically quantum properties for base-pair and dimer structures, are essential in positions 18-20 (Figure 3A; Figure 3C). In contrast, the seed region features further from the PAM sequence are specific to tetramer complexes. These differences suggest a structure-activity relationship, and may indicate variation in the mechanistic steps involving these regions. Looking to the multi-step CRISPR-Cas9 mechanism, we postulate that considering both DNA-DNA double helix unwinding and subsequent DNA-RNA binding are essential for interpreting these results.

This distinction can be seen when interpreting a positive correlation between the hydrogen bond stacking energy at position 18 of the target sequence (Figure 3A). Mechanistically, this indicates that position 18 is important for DNA-RNA binding. Once helix melting has been initiated at the target sequence’s 3’ end, the remaining sequence composition is less important for unwinding. Therefore, while lower hydrogen bond strength at position 20 is energetically preferable for DNA double helix melting, higher hydrogen bond strength at position 18 is important for strong DNA-RNA binding (Figure 3A).

In another view, a positive correlation between the HOMO-LUMO gap energy and cutting efficiency is observed at position 20. The HOMO-LUMO gap may capture conformational changes that are occurring during the initial integration of the CRISPR-Cas9 molecule. We note recent work identifying the “phosphate lock loop” in this interpretation. When the PAM sequence is identified and bound, the DNA “kinks” to enable DNA helix unwinding and permit DNA-RNA binding. These structural events are stabilized by a phosphate lock loop proximal to the PAM ^34–37^. While a high HOMO-LUMO gap at this region may describe a change in molecular stability, the weaker hydrogen bond may relate to the DNA double helix unwinding that follows.

A further discovery was the variation in influential properties for sgRNA efficiency across species. The novel quantum chemical property feature set is transferable across species because of its construction from simple nucleotide sequences. While the model does not provide a comprehensive view of nucleotide binding and interaction in complex genomes, it does provide a structural grounding for mechanistic interpretations that cannot be captured with traditional binary encodings. Species-tailored iRF models generated utilizing *E. coli* and *H. sapien* data exemplify this increase in interpretability. Moreover, the *H. sapien* model provides sufficient predictive power while allowing for feature engineering insight not currently available with top predictive models in the field due to their deep learning focus ^38^. This model performance points to the beneficial integration of quantum chemical properties, not only for interpretation but for sgRNA efficiency prediction.

While this novel feature set was shown to be of high importance across species, the specific quantum chemical properties differed. This highlights its use for understanding complex mechanisms across diverse species sets and supports critiques that current organism-tailored models are not applicable across species. This was further emphasized by the very low performance of the *E. coli* trained model as a predictor of *H. sapien* sgRNA efficiency.

This work established a novel feature set, with quantum chemical tensors, that advances the mechanistic interpretation and predictive accuracy of the model and will become a resource for continued work in the field. Initial insights into essential variables in the understanding of the CRISPR-Cas9 mechanism have been identified through the use of feature engineering techniques. These advances provide avenues for improving CRISPR-Cas9 sgRNA generation, identify points of interest for experimental assessment of the CRISPR-Cas9 mechanism, provide insights into species variation of CRISPR-Cas9, and provide methods for predictive model enhancement.

## Methods

### Datasets

#### E. coli

A publicly accessible *Escherichia coli* dataset published by ^28^ was utilized. Briefly, this dataset contains 55,670 unique sgRNAs that are profiled by co-expressing a genome-scale library with a pooled screening strategy. The data was established for three separate conditions including Cas9 (*Streptococcus pyogenes*), eSpCas9, and Cas9 (ΔrecA). The eSpCas9 is a Cas9 that has been reengineered for improved specificity and the Cas9 (ΔrecA) was developed by knockout of recA blocking DSBs repair. The dataset contains both sgRNA sequence and empirical CRISPR-Cas9 efficiency scores for each of the respective guides. The cutting efficiency scores were calculated by taking the log2 of the selected read count to the control read count. We focused on the Cas9 dataset for analyses within this manuscript.

#### H. sapien

A publicly accessible *H. sapien* dataset published by ^10^ was utilized. This dataset contains 1,278 unique sgRNAs based on an A375 viability analysis. The cutting efficiency was determined in the same manner as described above with the log2 fold change calculated relative to the change in abundance during a two week growth period. Additionally, a larger curated dataset by ^33^ which contains four publicly accessible human experimental sgRNA efficiency datasets ^11,19,39^ including multiple cell lines (HCT116, HEK293T, HELA, and HL60). The cutting efficiency value was defined as the log fold change in the measured knockout efficacy.

#### Multi-species

The multi-species model included sgRNA efficiency data from all previously described datasets. When model training occurs on multiple species datasets, all data is normalized on a scale of 0 to 1 and combined into a single matrix of sgRNA and cutting efficiency scores for model input. To eliminate species bias due to sample size, a consistent subsampling of 15,000 sgRNAs was utilized from *E. coli* and *H. sapien*, and the species was encoded as a binary feature.

### Feature Matrix

#### Quantum Chemical Properties ^29^

Quantum chemical properties provide unique insights into the factors influencing sgRNA efficiency in CRISPR-Cas9 systems. Duplexes of DNA-DNA, and DNA-RNA were modeled to assess these factors. The analysis includedincludedThese included e quantum chemical properties of individual bases; base-pairs; and base-pair dimers, trimers, and tetramers. In this way, a fourteen-Ångstrom (four base-pair) cut-off distance was invoked for long-range quantum interactions in the sgRNA. Additionally, a new sliding-window approach for the nucleotide base positions was developed for sgRNA interactions with the target DNA. Base-pair interactions were encoded into blocks, which subdivided the twenty-nucleotide sgRNA. In this approach, five ranges of interactions were assessed, from intramolecular to intermolecular.

HOMO-LUMO gap has been described as a signpost for a molecule’s kinetic stability ^40^. It describes the energetics of allowed electron transitions, and the likelihood of processes involving electron mobility. Structurally, the H-L gap reflects a molecule’s landscape of phase dependence for wave function interactions—both constructive and destructive—that originate covalent molecular interactions ^41^. Meanwhile, hydrogen bonding is a contextual property. It describes an energetic preference for arrangements of molecules in relation to each other. Hydrogen bonding directs non-covalent interactions between molecules, playing roles in thermodynamic stability and the energetics of protein folding, as two examples ^42^. Stacking interactions are similarly contextual interactions, and occur between aromatic rings. Stacking interactions range from π-π interactions within the rings—of the overlapping *p*-orbital electron density—to steric repulsions from exocyclic groups, which are implicated in DNA twisting ^43^. In this way, hydrogen bonding and stacking interactions differ in the chemical species that participate. Whereas hydrogen bonding interactions occur between hydrogen and a hydrogen bond acceptor, stacking interactions occur between aromatic species. Quantifying the energetics of these interactions complements a detailed feature set for machine learning models for sgRNA efficiency prediction.

The density-functional-based tight binding method (DFTB) is a powerful approach for large-scale atomistic simulations and calculating quantum properties. This work uses the DFTB3/3ob parameter set (third order parametrization for biological and organic systems). Calculations with the DFTB3/3ob parameter set yield excellent molecular geometries, which compare favorably with more resource-intensive methods. For example, DFTB3-3OB structures exhibit maximum absolute deviations of 0.045 Ångstroms from MP2/6-31G(d) methods (Second order Møller– Plesset perturbation theory with six-primitive split valence polarized Pople basis; ^44^).

##### Initial Coordinates

Nucleotide coordinates were collected from PubChem ^45^. B-DNA base-pairs were extracted from crystal structure data ^46^; PDBID: 167D). RNA hybrids and DNA-RNA hybrids were prepared by sterics-driven structure overlay in Biovia Discovery Studio software (Dassault Systèmes, S.E.). The nucleotide base and base-pair geometries were optimized through a gradient descent method with the simulation procedure described below. After optimization, each base-pair was aligned with the xy-plane in Open-Pymol software (Schrödinger, Inc.). In analogy to Gil *et al*. ^47^, each base-pair was then translated such that its centroid was the origin of the coordinates. To complete the unambiguous set of transformations, the pyrimidine carboxyl groups provided a final constraint. For this, the thymine carbonyl bond and cytosine carbonyl carbon were rotated to be normal to each other.

##### K-mer Construction

For all constructs, base-pairs were stacked at a distance of 3.5 Å along the z-axis, and rotated 36 degrees about their centroids (the origin). Structures were prepared with scripts executed in Open-Pymol. All non-chimeric single-strand combinations were assessed. In total, four bases, four base-pairs, 16 base-pair dimers, 64 base-pair trimers, and 256 base-pair tetramers were evaluated. Compiled k-mers were assessed by single-point energy calculations using the simulation procedure described below.

##### Simulation

Calculations were carried out at the DFTB3-D3(BJ)/3ob level of theory with Grimme’s D3(BJ) dispersion correction ^48,49^. Dispersion corrections were included to capture non-covalent interactions, resolving van der Waals and London dispersion forces in detail. Grimme’s dispersion correction was selected to describe medium and short-range dispersion effects ^50^. Additionally, a “COSMO” model was used with water as a solvent (<u>co</u>nductor-like <u>s</u>creening <u>mo</u>del). This model approximated solvent interactions, and contextualized the geometries and energy calculations to a water environment. Total system energy, HOMO-LUMO gap, and other quantum tensors were compiled for assessment in an Iterative Random Forest model.

#### Positional Encoding

Matrix generation incorporated extraction of several isolated sets of features. Position-independent and position-dependent positional encoding of the 20bp sgRNA was done as described by Doench et al., 2014. Briefly, position-independent features were determined by the count of nucleotides within the 20bp sequence both as a single base (A/C/T/G) and as paired bases (AA/AC/AG/etc.). Position-dependent features were represented using a binary variable (0 or 1) to encode the position-dependent single or paired base. Therefore, each position is encoded with a binary value for each of the four possible bases (A/C/T/G) with the nucleotide present at that position encoded as a value of one. Paired bases are further encoded with a binary value for each of the 16 possible base pair combinations. Additionally, we encoded the PAM (NGG) sequence by incorporating position-independent encoding of the N nucleotide. The combination of positional encoding approaches resulted in 384 features for each sgRNA assessed.

Further positional encoding was conducted on a kmer basis in order to incorporate larger scale combinations of nucleotides where the surrounding patterns influence sgRNA binding and efficiency. This was done through a stepwise integration of additional nucleotides as described above in a position-dependent manner including nucleotides in groups of two to five. The binary matrix includes the positional encoding using a sliding window so that each position from 1 to 20 minus the kmer length is encoded.

#### “Raw”features

Several raw value features were determined including GC content (ratio from 0-1 representing the proportion of the sgRNA sequence that is composed of GC), temperature of melting of the DNA duplex (calculated by the Watson-Crick formula of Tm(°C) = 64.9 + 41 * (nG+nC-16.4)/(nA+nT+nG+nC)), minimum free energy as a representation of RNA structure (calculated with ViennaRNA; ^51^), distance of the target sequence to the closest downstream PAM (utilizing the known genome assembly this was determined by the number of bases between position 20 of the sgRNA and the nearest NGG), and location relative to the target gene (represented by TSS, TTS, and quartiles of gene sequence (Q1-Q4)). These calculated values resulted in an additional 5 features for each sgRNA assessed.

### Iterative Random Forest Model

Random forest (RF) is a non-linear regression model which incorporates an ensemble of decision trees that trace the algorithm’s decision process. Iterative Random Forest (iRF) expands on the Random Forest method and is described in ^52,53^. Briefly, iRF is an advanced form of RF that implements a boosting and feature culling process based on the feature importance values from the previous iteration’s random forest to further iterate and amplify the features that repeatedly indicate high predictive capacity. It adds an iterative boosting process, producing a similar effect to Lasso in a linear model framework. In iRF, a Random Forest is created where features are unweighted and randomly sampled, at any given node in the decision trees, with equal probability. This process generates feature importance scores that are used to weight features in the next forest. This iterative method provides an amplification effect, increasing the chance that important features are evaluated at any given node ^52,53^.

For this study, the process of weighting and creating a new Random Forest is repeated 10 times with 1000 trees and incorporates a 5-fold cross validation. For each run, the data is separated into an 80/20 training/test split where 80% of the data is used for training the model and the remaining data (not utilized in model training) is used for testing. Each feature is ranked by its importance in the model in the tree building step and the direction of impact is determined based on the correlation of feature value and cutting efficiency value. Specifically, the feature matrix described above (incorporating positional encoding of the nucleotide composition of the sgRNA through a one-hot binary method and quantitative quantum chemical property values) is utilized to predict the cutting efficiency of the sgRNA. Multiple iRF models were run utilizing different feature sets to best understand the features of top importance and predictive capabilities; details on these models are in Table 1 below.

### Advanced Feature Engineering Metrics

In combination with iterative Random Forest, in-house scripts for advanced interpretation of machine learning output were utilized:

#### Random intersection trees (RIT)

is a method using binary predictor variables to identify interactions between features in a model. In short, RIT starts with a high level interaction that includes all variables in the matrix and then gradually removes variables as they fail to appear in randomly chosen observations of a specified class of interest ^53,54^. The algorithm works by assessing the node-split forest paths from iRF to find features that occur consecutively along the path more than would be expected by chance. The result of RIT is a set of interactions that have been retained with high probability, potentially in a non-linear manner, and are therefore considered informative to the model as a whole when joint.

For the analyses described we utilize internal R and python scripts specifically designed to deal with the extensive and iterative tree-based decision process of iRF and expand upon traditional RIT methods for enhanced feature engineering and model comparison. The resulting metrics for feature interpretation include normalized importance scores (in order to compare importances across models), feature effect scores (that show the magnitude and direction of the feature on the model), and number of samples captured as well as RIT (prevalence of set in the model), RIT adjusted (difference of prevalence of set in the model from the expected prevalence of the set), and set importance for identification and characterization of interacting features.

## Supporting information

Supplemental Results

## Code Availability

Github for iRF [https://github.com/jailGroup/RangerBasediRF] Github for feature matrix [https://github.com/nosha003/sgRNA_iRF]

## Acknowledgements/Funding Sources

This work was funded by the Secure Ecosystem & Engineering Design Science Focus Area is sponsored by the Genomic Science Program, U.S Department of Energy, Office of Science, Biological and Environmental Research, under FWP ERKPA17. This work was also funded by the Center for Bioenergy Innovation, a DOE Bioenergy Research Center supported by the Office of Biological and Environmental Research in the DOE Office of Science. Oak Ridge National Laboratory is managed by UT-Battelle, LLC for the U.S. Department of Energy under contract no. DE-AC05-00OR45678. This program is supported by the U. S. Department of Energy, Office of Science, through the Genomic Science Program, Office of Biological and Environmental Research, under FWP ERKP123. This work was also funded by a DOE Office of Energy Efficiency and Renewable Energy workforce development program. This research used resources of the Oak Ridge Leadership Computing Facility at the Oak Ridge National Laboratory, which is supported by the Office of Science of the U.S. Department of Energy under Contract no. DE-AC05-00OR22725. This research used resources of the Compute and Data Environment for Science (CADES) at the Oak Ridge National Laboratory, which is supported by the Office of Science of the U.S. Department of Energy under Contract No. DE-AC05-00OR22725.

## Author Contributions

**Conceptualization**: DJ, DK, JMN; **Methodology**: JMN, DK, DJ; **Software**: JR (RIT, iRF); **Formal analysis**: JMN, TW; **Investigation**: JMN, DK; **Resources**: DJ; **Writing - Original Draft**: JMN, TW; **Writing - Review & Editing**: JMN, TW, AW, DK, DJ, SI, CE; **Visualization**: JMN, TW, EP; **Supervision**: DK, DJ; **Project administration**: JMN; **Funding acquisition**: DJ, DK; All authors provided critical commentary and contributed to editing the manuscript.

## CRediT Authorship Categories

**Conceptualization**: Ideas; formulation or evolution of overarching research goals and aims

**Methodology**: Development or design of methodology; creation of models

**Software**: Programming, software development; designing computer programs; implementation of the computer code and supporting algorithms; testing of existing code components

**Validation**: Verification, whether as a part of the activity or separate, of the overall replication/ reproducibility of results/experiments and other research outputs

**Formal analysis**: Application of statistical, mathematical, computational, or other formal techniques to analyze or synthesize study data

**Investigation**: Conducting a research and investigation process, specifically performing the experiments, or data/evidence collection

**Resources**: Provision of study materials, reagents, materials, patients, laboratory samples, animals, instrumentation, computing resources, or other analysis tools

**Data Curation**: Management activities to annotate (produce metadata), scrub data and maintain research data (including software code, where it is necessary for interpreting the data itself) for initial use and later reuse

**Writing - Original Draft**: Preparation, creation and/or presentation of the published work, specifically writing the initial draft (including substantive translation)

**Writing - Review & Editing**: Preparation, creation and/or presentation of the published work by those from the original research group, specifically critical review, commentary or revision – including pre- or post-publication stages

**Visualization**: Preparation, creation and/or presentation of the published work, specifically visualization/ data presentation

**Supervision**: Oversight and leadership responsibility for the research activity planning and execution, including mentorship external to the core team

**Project administration**: Management and coordination responsibility for the research activity planning and execution

**Funding acquisition**: Acquisition of the financial support for the project leading to this publication

---

**Figure S1:**
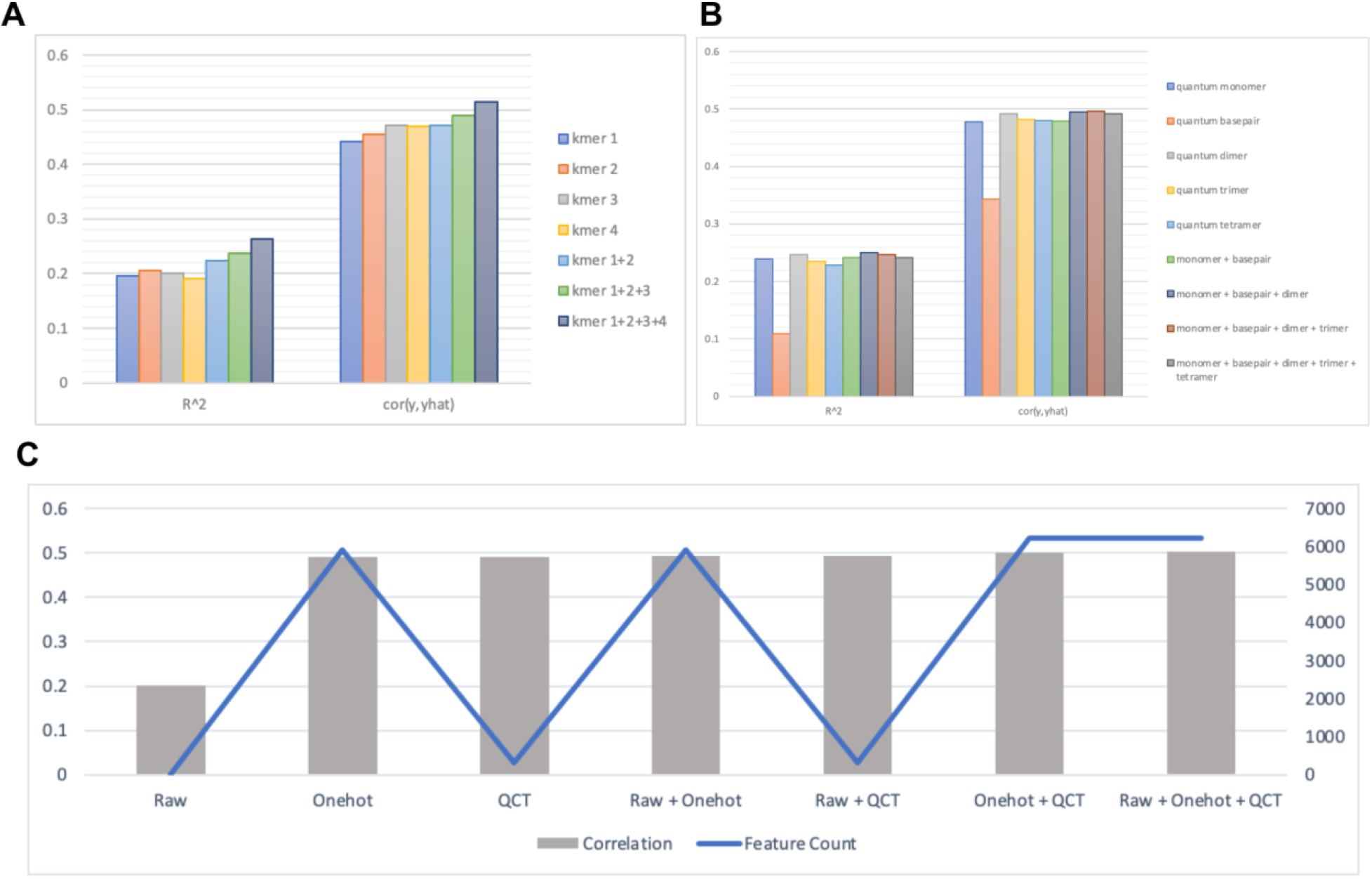
iRF metric output relative to feature set. Model output metrics including R squared and Pearson correlation of the predicted versus experimental cutting efficiency scores for A) k-mer integration of positional encoding features and B) k-mer integration of quantum property features. C) Bar plot of the Pearson correlation predictive metric from iRF output based on the feature matrix utilized and the corresponding number of features incorporated in that matrix (line plot) showing that the increase in matrix size is not the main contributor to high predictive metrics.

**Figure S2:**
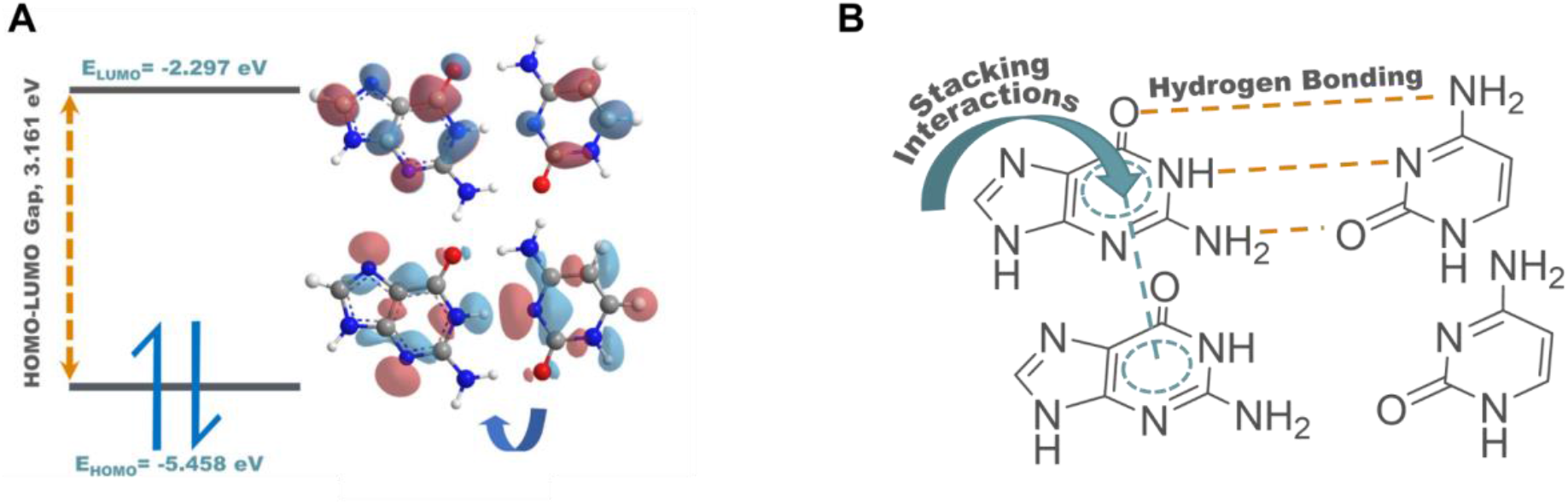
A) The identification of the highest and lowest occupied molecular orbitals for quantum property calculations (left). A visual depiction of the HOMO-LUMO gap based on the nucleotide organization of the sgRNA sequence (right). B) Schematic depiction of hydrogen bonding property and stacking interactions.

**Figure S3:**
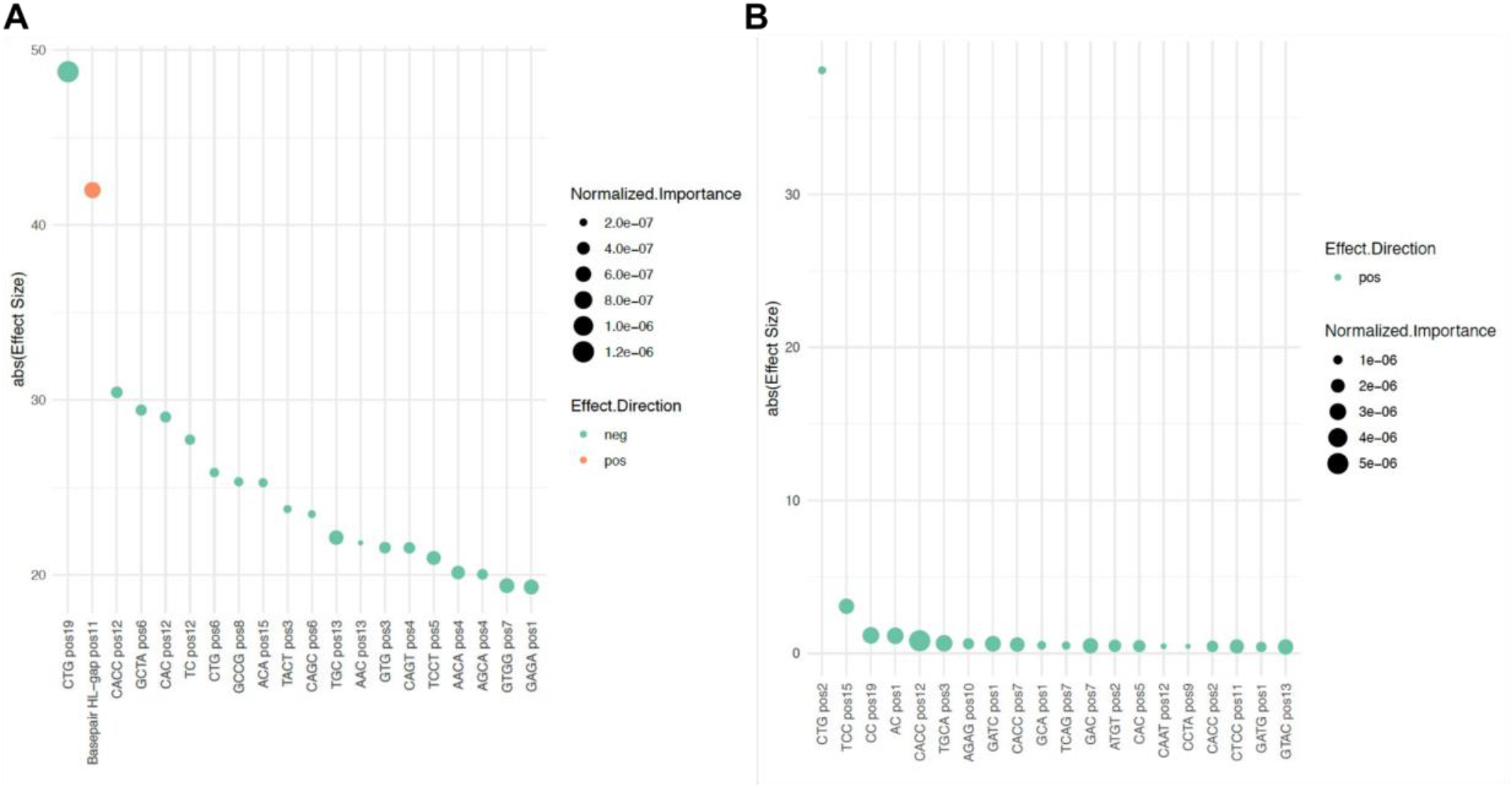
Effect size and direction for A) *E. coli* and B) *H. sapien*. Dot plots of top 20 features based on effect size, calculated as the average proportion of sgRNAs that were influenced by that feature based on the presence in decision trees of the iRF model. Additional information displayed includes the direction of effect (positive = orange, negative = green) and importance score (size of dot).

**Figure S4:**
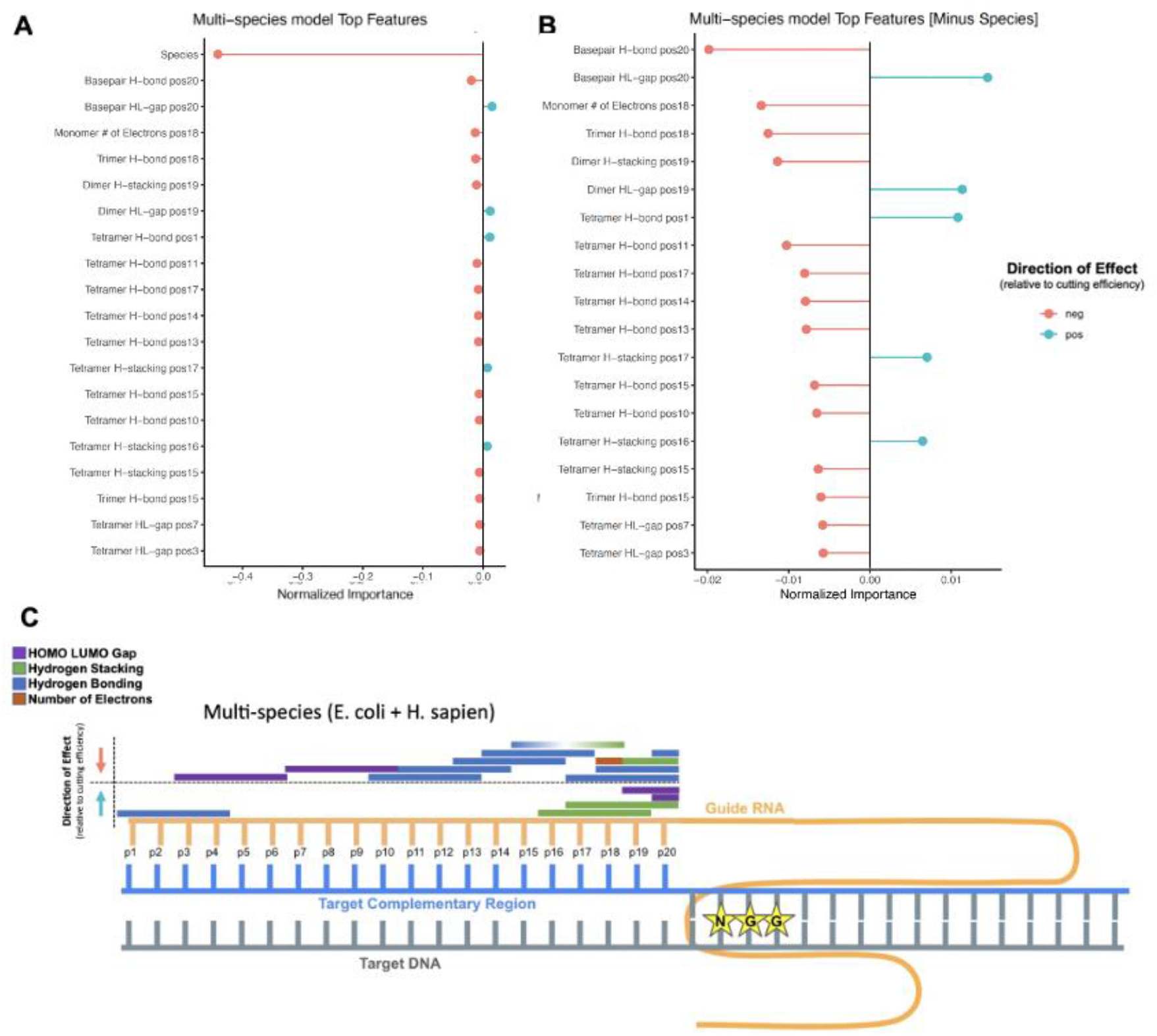
A) The top 20 features from the *E.coli* and *H.sapien* multi-species iRF model ranked by normalized importance score and colored by the direction of the effect as positively correlated with the cutting efficiency score (blue) or anti-correlated with cutting efficiency score (pink). B) The top 20 features minus the species feature to zoom in on the remaining more species-generalizable features. C) sgRNA-DNA interaction highlighting quantum chemical property features of top importance and their localization for positively correlated association and anticorrelated associations with cutting efficiency scores in the multi-species model. DNA strand represented in grey (target sequence) and blue (target complementary sequence), sgRNA shown in yellow, and PAM sequence displayed with NGG stars. Direction of feature effect is indicated with arrows, up (blue arrow) indicates a positively correlated relationship between the feature value and the cutting efficiency value. Feature bars indicate specific quantum properties (HL gap = purple, Stacking interactions = green, H-bonding = blue) and the length of the bar indicates the k-mer size.

## References

1. Naim, F. et al. Are the current gRNA ranking prediction algorithms useful for genome editing in plants? PLoS One 15, e0227994 (2020).

2. Doudna, J. A. & Charpentier, E. The new frontier of genome engineering with CRISPR-Cas9. Science 346, (2014).

3. Wu, X., Kriz, A. J. & Sharp, P. A. Target specificity of the CRISPR-Cas9 system. Quant Biol 2, 59–70 (2014).

4. Liu, G., Zhang, Y. & Zhang, T. Computational approaches for effective CRISPR guide RNA design and evaluation. Comput. Struct. Biotechnol. J. 18, 35–44 (2020).

5. Moreno-Mateos, M. A. et al. CRISPRscan: designing highly efficient sgRNAs for CRISPR-Cas9 targeting in vivo. Nat. Methods 12, 982–988 (2015).

6. Housden, B. E. et al. Identification of potential drug targets for tuberous sclerosis complex by synthetic screens combining CRISPR-based knockouts with RNAi. Sci. Signal. 8, rs9 (2015).

7. Labuhn, M. et al. Refined sgRNA efficacy prediction improves large-and small-scale CRISPR-Cas9 applications. Nucleic Acids Res. 46, 1375–1385 (2018).

8. Rahman, M. K. & Rahman, M. S. CRISPRpred: A flexible and efficient tool for sgRNAs on-target activity prediction in CRISPR/Cas9 systems. PLoS One 12, e0181943 (2017).

9. Tsai, S. Q. & Keith Joung, J. Defining and improving the genome-wide specificities of CRISPR–Cas9 nucleases. Nat. Rev. Genet. 17, 300–312 (2016).

10. Doench, J. G. et al. Rational design of highly active sgRNAs for CRISPR-Cas9-mediated gene inactivation. Nat. Biotechnol. 32, 1262–1267 (2014).

11. Wang, T., Wei, J. J., Sabatini, D. M. & Lander, E. S. Genetic screens in human cells using the CRISPR-Cas9 system. Science 343, 80–84 (2014).

12. Xu, H. et al. Sequence determinants of improved CRISPR sgRNA design. Genome Res. 25, 1147–1157 (2015).

13. Cong, L. et al. Multiplex genome engineering using CRISPR/Cas systems. Science 339, 819–823 (2013).

14. Liu, X. et al. Sequence features associated with the cleavage efficiency of CRISPR/Cas9 system. Scientific Reports vol. 6 (2016).

15. Mans, R. et al. CRISPR/Cas9: a molecular Swiss army knife for simultaneous introduction of multiple genetic modifications in Saccharomyces cerevisiae. FEMS Yeast Res. 15, (2015).

16. Bassett, A. R. & Liu, J. L. CRISPR/Cas9 and genome editing in Drosophila. J. Genet. Genomics (2014).

17. Liu, H. et al. CRISPR-P 2.0: An Improved CRISPR-Cas9 Tool for Genome Editing in Plants. Mol. Plant 10, 530–532 (2017).

18. Smith, J. D. et al. Quantitative CRISPR interference screens in yeast identify chemical-genetic interactions and new rules for guide RNA design. Genome Biol. 17, 45 (2016).

19. Doench, J. G. et al. Optimized sgRNA design to maximize activity and minimize off-target effects of CRISPR-Cas9. Nat. Biotechnol. 34, 184–191 (2016).

20. Abadi, S., Yan, W. X., Amar, D. & Mayrose, I. A machine learning approach for predicting *CRISPR-Cas9 cleavage efficiencies and patterns underlying its mechanism of action*. PLoS Comput. Biol. 13, e1005807 (2017).

21. Shibata, M. et al. Real-space and real-time dynamics of CRISPR-Cas9 visualized by high-speed atomic force microscopy. Nat. Commun. 8, 1430 (2017).

22. Horlbeck, M. A. et al. Nucleosomes impede Cas9 access to DNA in vivo and in vitro. Elife 5, (2016).

23. Gisler, S. et al. Multiplexed Cas9 targeting reveals genomic location effects and gRNA-based staggered breaks influencing mutation efficiency. Nat. Commun. 10, 1598 (2019).

24. Yarrington, R. M., Verma, S., Schwartz, S., Trautman, J. K. & Carroll, D. Nucleosomes inhibit target cleavage by CRISPR-Cas9 in vivo. Proc. Natl. Acad. Sci. U. S. A. 115, 9351–9358 (2018).

25. Chen, Y. et al. Using local chromatin structure to improve CRISPR/Cas9 efficiency in zebrafish. PLoS One 12, e0182528 (2017).

26. Jensen, K. T. et al. Chromatin accessibility and guide sequence secondary structure affect CRISPR-Cas9 gene editing efficiency. FEBS Letters vol. 591 1892–1901 (2017).

27. Lino, C. A., Harper, J. C., Carney, J. P. & Timlin, J. A. Delivering CRISPR: a review of the challenges and approaches. Drug Deliv. 25, 1234–1257 (2018).

28. Guo, J. et al. Improved sgRNA design in bacteria via genome-wide activity profiling. Nucleic Acids Res. 46, 7052–7069 (2018).

29. Gadiyaram, V., Vishveshwara, S. & Vishveshwara, S. From Quantum Chemistry to Networks in Biology: A Graph Spectral Approach to Protein Structure Analyses. J. Chem. Inf. Model. 59, 1715–1727 (2019).

30. McFadden, J. & Al-Khalili, J. The origins of quantum biology. Proc. Math. Phys. Eng. Sci. 474, 20180674 (2018).

31. Zhu, H. & Liang, C. CRISPR-DT: designing gRNAs for the CRISPR-Cpf1 system with improved target efficiency and specificity. bioRxiv 269910 (2018) doi:10.1101/269910.

32. Shah, R. D. & Meinshausen, N. Random intersection trees. J. Mach. Learn. Res. (2014).

33. Chuai, G. et al. DeepCRISPR: optimized CRISPR guide RNA design by deep learning. Genome Biol. 19, 80 (2018).

34. Palermo, G. et al. Key role of the REC lobe during CRISPR–Cas9 activation by ‘sensing’, ‘regulating’, and ‘locking’ the catalytic HNH domain. Quarterly Reviews of Biophysics vol. 51 (2018).

35. Raper, A. T., Stephenson, A. A. & Suo, Z. Functional Insights Revealed by the Kinetic Mechanism of CRISPR/Cas9. J. Am. Chem. Soc. 140, 2971–2984 (2018).

36. Nishimasu, H. et al. Crystal structure of Cas9 in complex with guide RNA and target DNA. Cell 156, 935–949 (2014).

37. Jiang, F. & Doudna, J. A. CRISPR–Cas9 Structures and Mechanisms. (2017) doi:10.1146/annurev-biophys-062215-010822.

38. Zhang, G., Dai, Z. & Dai, X. A Novel Hybrid CNN-SVR for CRISPR/Cas9 Guide RNA Activity Prediction. Front. Genet. 10, 1303 (2019).

39. Hart, T. et al. High-Resolution CRISPR Screens Reveal Fitness Genes and Genotype-Specific Cancer Liabilities. Cell 163, 1515–1526 (2015).

40. Aihara, J.-I. Reduced HOMO-LUMO Gap as an Index of Kinetic Stability for Polycyclic Aromatic Hydrocarbons. J. Phys. Chem. A 103, 7487–7495 (1999).

41. Levine, D. S. & Head-Gordon, M. Clarifying the quantum mechanical origin of the covalent chemical bond. Nat. Commun. 11, 4893 (2020).

42. Gao, J., Bosco, D. A., Powers, E. T. & Kelly, J. W. Localized thermodynamic coupling between hydrogen bonding and microenvironment polarity substantially stabilizes proteins. Nat. Struct. Mol. Biol. 16, 684–690 (2009).

43. Cooper, V. R. et al. Stacking interactions and the twist of DNA. J. Am. Chem. Soc. 130, 1304–1308 (2008).

44. Gaus, M., Goez, A. & Elstner, M. Parametrization and Benchmark of DFTB3 for Organic Molecules. J. Chem. Theory Comput. 9, 338–354 (2013).

45. Kim, S. et al. PubChem in 2021: new data content and improved web interfaces. Nucleic Acids Res. 49, D1388–D1395 (2021).

46. Goodsell, D. S., Kaczor-Grzeskowiak, M. & Dickerson, R. E. The crystal structure of C-C-A-T-T-A-A-T-G-G. Implications for bending of B-DNA at T-A steps. J. Mol. Biol. 239, 79–96 (1994).

47. Gil, A., Branchadell, V., Bertran, J. & Oliva, A. An Analysis of the Different Behavior of DNA and RNA through the Study of the Mutual Relationship between Stacking and Hydrogen Bonding. The Journal of Physical Chemistry B vol. 113 4907–4914 (2009).

48. Grimme, S., Antony, J., Ehrlich, S. & Krieg, H. A consistent and accurate ab initio parametrization of density functional dispersion correction (DFT-D) for the 94 elements H-Pu. J. Chem. Phys. 132, 154104 (2010).

49. Grimme, S., Ehrlich, S. & Goerigk, L. Effect of the damping function in dispersion corrected density functional theory. J. Comput. Chem. 32, 1456–1465 (2011).

50. Schröder, H., Creon, A. & Schwabe, T. Reformulation of the D3(Becke-Johnson) *Dispersion Correction without Resorting to Higher than C_6_ Dispersion Coefficients*. J. Chem. Theory Comput. 11, 3163–3170 (2015).

51. Lorenz, R. et al. ViennaRNA Package 2.0. Algorithms Mol. Biol. 6, 26 (2011).

52. Cliff, A. et al. A High-Performance Computing Implementation of Iterative Random Forest for the Creation of Predictive Expression Networks. Genes 10, (2019).

53. Basu, S., Kumbier, K., Brown, J. B. & Yu, B. Iterative random forests to discover predictive and stable high-order interactions. Proc. Natl. Acad. Sci. U. S. A. 115, 1943–1948 (2018).

54. Shah, R. D. & Meinshausen, N. Random Intersection Trees. arXiv [stat.ML] (2013).

