## Supplemental Results for "Quantum biological insights into CRISPR-Cas9 sgRNA efficiency from explainable-AI driven feature engineering"

**Supplemental Data:**
Supplemental Table 1: Quantum Chemical Properties
Supplemental Table 2: E.coli matrix
Supplemental Table 3: H.sapien matrix
Supplemental Table 4: E.coli iRF normalized feature importance scores

**Supplemental Results:**

***Assessing model accuracy across feature sets:***
A total of eight iRF models were generated (Table 1; Figure S1). Across the iRF models, each of three feature sets was assessed in isolation and in combination with the remaining features (Table 1; Figure 2A; Figure S1). The "raw" values feature set included sgRNA structure, GC content, melting temperature, distance to PAM, and distance to gene. This feature set exhibited minimal predictive power (Pearson correlation = 0.20; $R^2$=0.04) for cutting efficiency scores (Figure 2A; Table 1). The position-independent nucleotide sequence encoding feature set, comprising the A,C,T,G count in the complete sgRNA sequence, had a similarly minimal standalone predictive power. Crucially, and in contrast, other feature sets exhibited enhanced predictive power, including one-hot positional nucleotide encoding (Pearson correlation=0.49; $R^2$ = 0.26 ) and quantum chemical properties (Pearson correlation=0.49; $R^2$ =0.24) (Figure 2A; Table 1). Additional improvements are observed when these features encompass k-mers beyond single and paired bases–to include base-pair dimer, trimer, and tetramer encodings (Figure S1A-B). The model accuracy increased to $R^2$=0.26 when these quantum chemical descriptors were considered with the "raw" feature set. Notably, these differences in predictive power do not follow the trend of matrix size as the Pearson correlation is agnostic to increases in data input. This behavior means that the types and breadth of information held within the feature sets are imperative for predictive power (Figure S1C). Each feature's contribution to the model's overall predictive power can also be assessed.

Surprisingly, comparing the predictive accuracy ($R^2$) across models, the five most important features captured only a small portion of the full model's accuracy (Figure 2D; FIgure S1C). This highlights the importance of combined and cumulative impacts from features which alone had small effects. Meanwhile, comparing Pearson correlations, the top 50 features retained most of the full model's predictive power. Minimal increases in prediction accuracy were observed in models beyond 500 features. (Figure 2D; Figure S1C). Even the top 1000 features do not reach the full predictive power of the original 6,232 features; however, they capture a majority of that capacity at lower computational cost. This reduced model provides a tractable system for investigating features' importance for sgRNA efficiency. Overall, These varied metrics exemplify the interpretability of our iRF model for CRISPR-Cas9 efficiency. Further, novel addition of quantum chemical properties provides insights for interpreting the complex biological mechanisms at play in CRISPR-based genome editing.

***A determination of individual feature effect***
The XAI approach provides powerful metrics to extend the analysis beyond the direction of a feature's relationship with cutting efficiency, to also assess the effect size. Effect size captures the rate at which the cutting score (the Y-vector) changes in response to a feature. The HOMO-LUMO gap at base pair 11 had the steepest effect size (Figure S3). Other top effect sizes include positional encoding features (Figure S3). This metric also lends interpretation to the model. The one-hot

encoding features capture the position-dependent encoding of the target sequence's 5' end. This region includes highly ranked dimer, trimer, and tetramer encoding of positions 1, 3, 6, and 11. These features all exhibited negative correlations, with the trimer and tetramer-defined base pair sequences having the greatest effects (Figure S3). Structurally, this suggests that specific nucleotides at these positions in the target DNA double helix negatively impact the cutting efficiency Interestingly, this contrasts with the most-important quantum chemical properties, which are localized to the target sequence's 3' end. Furthermore, all features with high effect size were positively correlated with cutting efficiency scores. These varied metrics allow for mechanistic interpretation and sgRNA feature engineering.

***Generating a multi-species sgRNA model***

Finally, a multi-species model was trained utilizing both the *E.coli* and *H.sapien* datasets described [10,28,33]. Cutting efficiency scores were normalized and a random sample of 15,000 sgRNAs from each dataset was used to eliminate bias of power from single-species contributions. The multi-species model was tested with a mixed *E.coli* and *H.sapien* dataset–not included in the training of the model–which showed the highest model accuracy of all iRF models generated, with an $R^2$ of 0.486. The multi-species test data also demonstrated a high predictive capability, with a Pearson correlation of 0.697. The test data to assess the predictive ability for each species. Both the *E.coli* and the *H.sapien* data had comparable Pearson correlations with the multi-species model and the species-tailored models (*E. coli:* species-tailored model=0.502, multi-species model=0.504) (*H. sapien:* species-tailored model=0.496, multi-species model=0.491).

The species was also encoded as a binary feature in the multi-species model and was identified as the top most important feature in the model. While this is essential to note, the specific training data used is compiled from multiple studies with varied methods. A caveat to this compiled dataset is the distinct skews in distribution of cutting efficiencies between the *E. coli* and *H. sapien* datasets. Normalizing the distribution of cutting efficiency scores across studies is not straightforward, and uncertainties influence the quantitative prediction values of the model. The top most important features identified are consistent with a mixed model; and highlight seed region quantum chemical properties, along with properties discussed for the *E.coli* and *H.sapien*-specific models (Figure S4).
